# p63 cooperates with CTCF to modulate chromatin architecture in skin keratinocytes

**DOI:** 10.1101/525667

**Authors:** Jieqiong Qu, Guoqiang Yi, Huiqing Zhou

## Abstract

The transcription factor p63 regulates epidermal genes and the enhancer landscape in skin keratinocytes. Its molecular function in controlling the chromatin structure is however not yet completely understood. Here we integrated multi-omics profiles, including the transcriptome, transcription factor DNA-binding and chromatin accessibility, in skin keratinocytes isolated from EEC syndrome patients carrying p63 mutations, to examine the role of p63 in shaping the chromatin architecture. We found decreased chromatin accessibility in p63-and CTCF-bound open chromatin regions that potentially contributed to gene deregulation in mutant keratinocytes. Cooperation of p63 and CTCF seemed to assist chromatin interactions between p63-bound enhancers and gene promoters in skin keratinocytes. Our study suggests an intriguing model where cell type-specific transcription factors such as p63 cooperate with the genome organizer CTCF in the three-dimensional chromatin space to regulate the transcription program important for the proper cell identity.

## Introduction

Skin development and homeostasis requires tightly regulated epidermal keratinocyte proliferation, differentiation and apoptosis, and these processes are governed by cooperation of cis-regulatory elements, transcription factors (TFs), chromatin accessibility as well as higher-order chromatin organisation [1-5]. The TF p63 is a key regulator of epidermal development. At the molecular level, p63 regulates a large number of genes, for example, p21 (*CDKN1A*) in cell cycle arrest [6], Fras1 in maintaining basement membrane integrity [7] and key genes such as keratins, filaggrin, and loricrin required for epidermal morphogenesis and differentiation [1, 8]. Furthermore, p63 directly regulates chromatin factors, Satb1, Lsh, and Brg1, to control the chromatin remodelling during epidermis development [9-11]. Recent studies showed that p63 exerts a crucial role in establishing the enhancer landscape [4, 5, 12, 13]. Through active enhancers, p63 cooperates with its co-regulating TFs to modulate local transcriptional program required for epidermal homeostasis [4].

In human, heterozygous mutations of *TP63* encoding p63 cause a spectrum of ectoderm-related disorders [14]. For example, Ectrodactyly-Ectodermal Dysplasia-Cleft Lip/Palate (EEC) syndrome is caused by point mutations located in the p63 DNA-binding domain, and manifests ectodermal dysplasia with defects in the epidermis and epidermal related appendages, limb malformation and cleft lip/palate. Five hotspot mutations affecting amino acids, R204, R227, R279, R280 and R304, cover approximately 90% of the all EEC syndrome cases [15]. Our previous study showed that mutant p63 resulted in a genome-wide redistribution of enhancers in keratinocytes established from EEC patients [13]. Consistently, the gene network analysis identified a significant co-expression gene module of ‘nucleosome assembly’, implying a less-organized chromatin structure in EEC syndrome keratinocytes [13]. How mutant p63 affects the chromatin structure is however not yet clear.

In this study, we characterized the chromatin accessibility using ATAC-seq in keratinocytes established from EEC patients carrying p63 mutations, in comparison with control keratinocytes. A clear difference in chromatin accessibility that correlated with the transcriptional dynamics was detected. Unexpectedly, strong enrichment of CTCF binding siteswere observed at control-specific open chromatin regions. By combining published promoter Capture Hi-C seq data, we found that CTCF and p63 were cooperatively involved in DNA loops to regulate epidermal genes. Our findings provide new insights into the coordinated regulatory role of CTCF and p63 in chromatin interactions in epidermal keratinocytes.

## Results

### Differential chromatin accessibility between control and p63 mutant keratinocytes

We performed Assay for Transposase-Accessible Chromatin with high-throughput sequencing (ATAC-seq) to characterize the accessible genome as well as the nucleosome position in both control and p63 mutant keratinocytes at the proliferation stage. Two replicas of ATAC-seq analyses showed high correlation (Figure 1A), indicating high reproducibility. The principle component analysis (PCA) plot based on the chromatin accessibility displayed a clear separation between control and p63 mutant keratinocytes (Figure 1B), which is highly consistent with results from gene expression profiles (Figure 1B). The PCA analysis also showed that the genome accessibility of the two p63 mutant lines (R204W and R304W) are similar, as they are close on the PC1 axis that represents major variations. As the goal of this work is to examine the difference between control and EEC mutant lines, these two mutant lines were considered as one group, termed as p63 mutant keratinocytes, in the following analyses.

**Figure 1.**
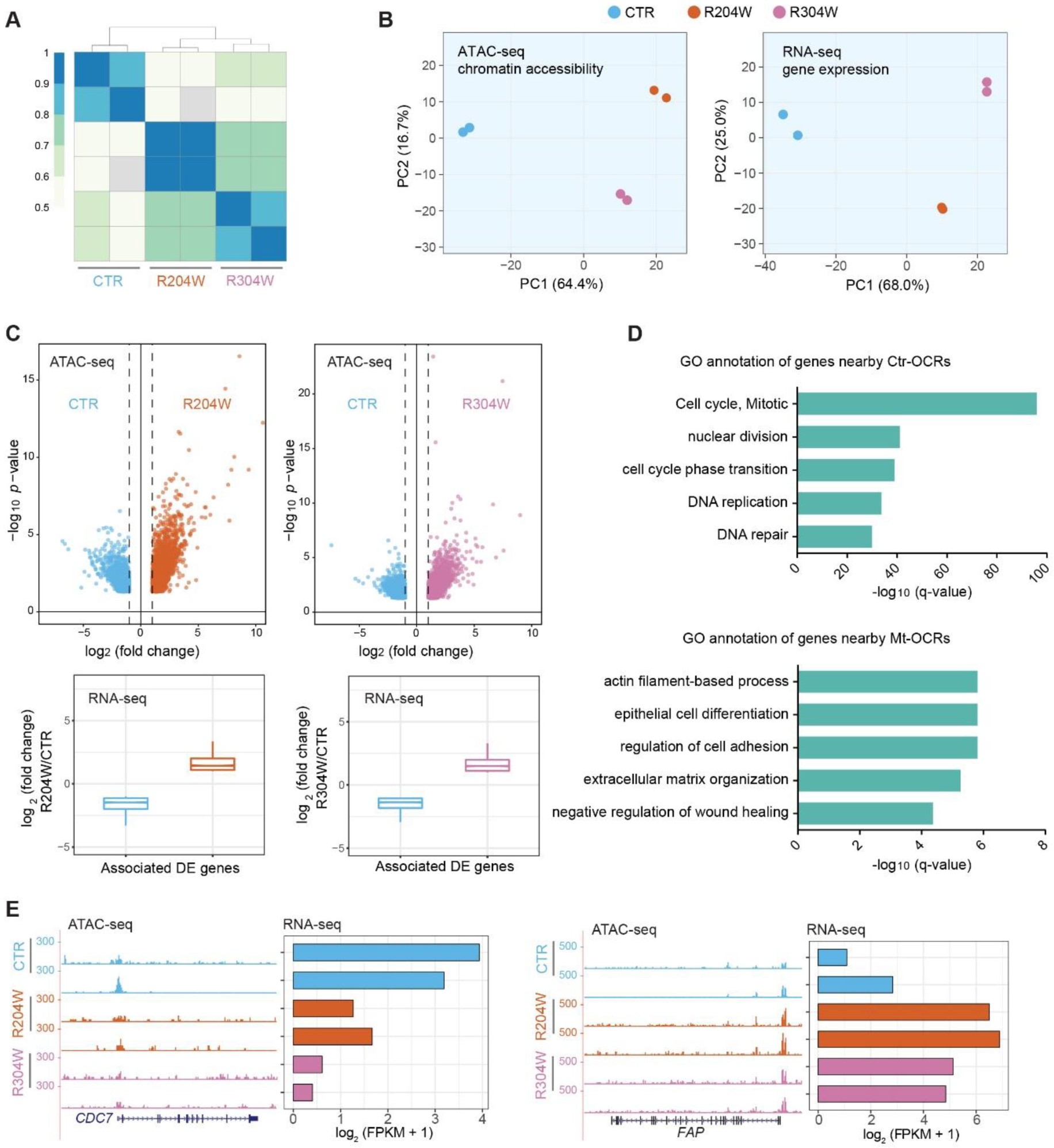
Differential chromatin accessibility in p63 mutant keratinocytes. A. A heatmap of sample correlation matrix showing high similarities between duplicates and dissimilarities between control (CTR) and p63 mutant keratinocytes (R204W and R304W). B. PCA plots of ATAC-seq and RNA-seq of control and p63 mutant keratinocytes at the proliferation stage. C. Upper panel, volcano plots of ATAC-seq comparisons between control and p63 mutant keratinocytes. The x axis shows the log2 fold change of reads detected at the open chromatin regions (OCRs) and the y axis shows −log10 (p value). Lower panel, the transcriptional changes of differentially expressed genes associated with differential OCRs indicated in the volcano plots. D. GO annotation of differentially expressed genes nearby Ctr-OCRs and Mt-OCRs. E. Examples of Ctr-OCRs and their associated DE gene, *CDC7*, as well as Mt-OCRs and their associated DE gene, *FAP*. ATAC-seq tracks are shown in the RPKM scale.

Subsequently we identified the differential open chromatin regions (OCRs) marked by ATAC-seq signals between control and p63 mutant keratinocytes. In total, there were 2,492 open chromatin regions that showed higher signal in control keratinocytes, termed as control-specific open chromatin regions (Ctr-OCRs); in parallel, there were 3,716 regions that showed higher signal in mutant keratinocytes, termed as mutant-specific open chromatin regions (Mt-OCRs) (Figure 1C). As expected, we found that differential OCRs were positively correlated with gene expression, when they were assigned to the nearest differentially expressed (DE) genes (Figure 1C). Genes associated with Ctr-OCRs were mainly involved in the regulation of ‘cell cycle’ (Figure 1D), e.g. *CDC7* (Figure 1E), while genes nearby Mt-OCRs were mainly enriched in ‘actin filament-based process’ (Figure 1D), e.g. *FAP* (Figure 1E).

### p63 and CTCF occupancy at Ctr-OCRs

To identify the potential TFs involved in these differential OCRs, we first performed a comparative motif analysis between control and p63 mutant keratinocytes using HOMER. Among the Ctr-OCRs, we found that p63 motif is most enriched (Figure 2A), which is consistent with our previous finding that loss of p63 binding resulted in decreased enhancers in p63 mutant keratinocytes [13].

**Figure 2.**
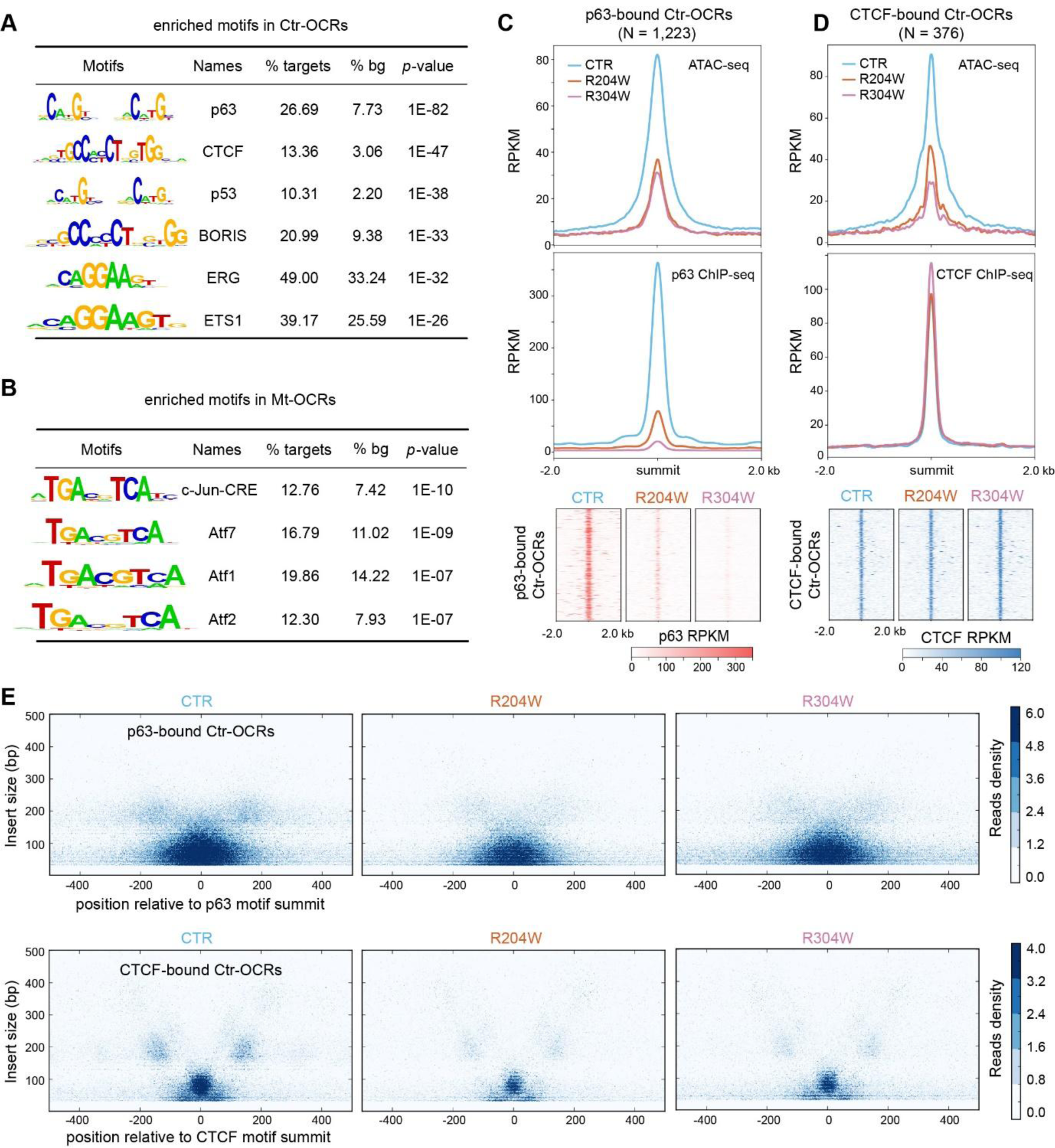
Decreased chromatin accessibility and altered nucleosome organization at p63-bound OCRs and CTCF-bound OCRs in p63 mutant keratinocytes. A. Top enriched motifs in Ctr-OCRs. 2,492 Ctr-OCRs (targets) were used in this analysis with Mt-OCRs as background (bg). B. Top enriched motifs in Mt-OCRs. 3,716 Mt-OCRs (targets) were used in this analysis with Ctr-OCRs as background (bg). C. Bandplots (top) and heatmaps (bottom) showing the quantification of ATAC-seq and p63 ChIP-seq signals at the 1,223 p63-bound Ctr-OCRs. D. Bandplots (top) and heatmaps (bottom) showing the quantification of ATAC-seq and CTCF ChIP-seq signals at the 376 CTCF-bound Ctr-OCRs. E. V-plot of ATAC-seq reads showing the read length and distance from either p63 motif (top) or CTCF motif (bottom) in control and p63 mutant keratinocytes. The short fragments (<150 bp) were indicative of nucleosome free DNA.

Unexpectedly, the motif of CTCF that is a well-known genome organizer [16] was identified as the 2^nd^ highly enriched motif in the Ctr-OCRs. Among the Mt-OCRs, AP-1 motif family was most enriched (Figure 2B), consistent with findings on gained enhancers in p63 mutant keratinocytes [13].

To validate our motif analysis result and explore the role of p63 in controlling Ctr-OCRs, we used published p63 ChIP-seq data from our previous work [13]. In control keratinocytes, a total of 1,223 p63 binding sites (BSs) were covered by the 2,492 Ctr-OCRs, showing a significant overlap (*p* < 0.001). When ATAC-seq and p63 ChIP-seq reads were mapped to the 1,223 p63-bound Ctr-OCRs, we found a decrease of both chromatin accessibility and p63 binding signals at the p63-bound Ctr-OCRs in p63 mutant keratinocytes (Figure 2C).

To confirm the role of CTCF in Ctr-OCRs, we performed CTCF ChIP-seq in both control and p63 mutant keratinocytes. In total, we found 12,116 CTCF BSs, among which only 363 were differential in CTCF binding between control and p63 mutant keratinocytes (*p* < 0.05), indicating that CTCF binding is rather stable. In addition, among all putative CTCF BSs, we found 376 overlapped with the 2,492 Ctr-OCRs, which is statistically significant (*p* < 0.001). We then mapped ATAC-seq and CTCF ChIP-seq reads to the 376 CTCF-bound Ctr-OCRs. As expected, decrease of ATAC-seq signals was observed at the CTCF-bound Ctr-OCRs in p63 mutant keratinocytes. However, unexpectedly, there was no significant change in CTCF binding signals at the CTCF-bound Ctr-OCRs in p63 mutant keratinocytes, as compared to the control keratinocytes (Figure 2D). Another expected finding was that there is few overlap (N=435) out of 45,350 p63 BSs and 12,116 CTCF BSs (Supplementary Figure 1A), although both motifs were identified in Ctr-OCRs. Consistently, there was no visible CTCF binding signal at all p63 BSs, and vice versa, no visible p63 binding at the CTCF BSs (Supplementary Figure 1B). Consistently, we also observed decreased p63 binding and largely unchanged CTCF binding signal in p63 mutant keratinocytes at the co-localized binding sites (Supplementary Figure 1C).

Subsequently, we analyzed the nucleosome positioning at p63-bound and CTCF-bound Ctr-OCRs by mapping the distribution of ATAC-seq fragments centered on either p63 motif or CTCF motif (Figure 2E). In control keratinocytes, we found an enrichment of short fragments (< 150 bp) surrounding both p63 motif and CTCF motif. The enrichment of such short reads clearly decreased in p63 mutant keratinocytes (Figure 2E and Supplementary Figure 2). We also observed an enrichment of fragments that had lengths slightly shorter than 200 bp which represent the stable flanking nucleosomes in control keratinocytes. However, in p63 mutant keratinocytes, the approximate 200 bp length-fragments seemed to be less present (Supplementary Figure 2), indicating an altered nucleosome organization.

Taken together, we observed decreased accessibility at p63-bound and CTCF-bound Ctr-OCRs in p63 mutant keratinocytes when compared to control keratinocytes. Notably, there was a clear decrease of p63 binding at the p63-bound Ctr-OCRs whereas CTCF binding did not seem to be affected at the CTCF-bound Ctr-OCRs in p63 mutant keratinocytes.

### Characterization of p63-bound OCRs and CTCF-bound OCRs

We reasoned that the chromatin states and the genomic locations of these Ctr-OCRs might be informative to understand the difference between p63-bound and CTCF-bound Ctr-OCRs. To this end, we firstly analyzed the chromatin states (CSs) of all CTCF and p63 BSs with the combination of six histone marks from Roadmap project [17] (Figure 3A). We found that p63 bound to both promoters and enhancers marked by H3K27ac and H3K4me1, whereas CTCF was more enriched at promoters marked by H3K4me3 (Figure 3A and Supplementary Figure 3). We then performed analyses on the p63-and CTCF-bound Ctr-OCRs (Figure 3B). When comparing to all Ctr-OCRs, p63-bound Ctr-OCRs were overrepresented at enhancers (CSs 8, 9, and 11) while the majority of CTCF-bound Ctr-OCRs showed overrepresentation at promoter proximal regions (CSs 2 and 4, Figure 3B), consistent with all p63 and CTCF BSs.

**Figure 3.**
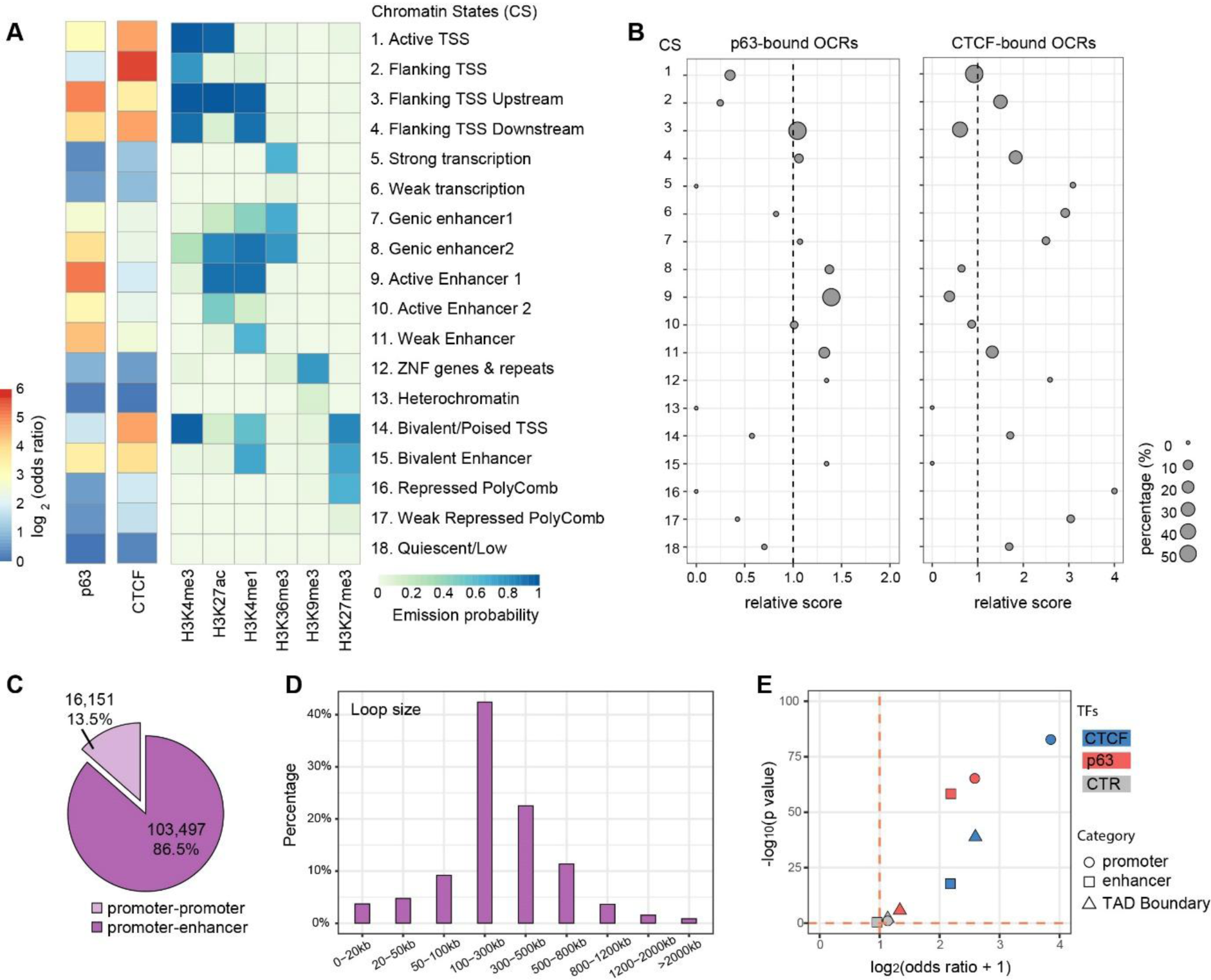
Chromatin state and locations of p63- and CTCF-bound OCRs. A. Fold enrichment/depletion of chromatin states (CSs) in p63 BSs and CTCF BSs compared to the randomized sequences. The 18 CSs were defined using ENCODE data [17]. B. Relative enrichment (> 1)/depletion (< 1) of CSs in p63-bound Ctr-OCRs and CTCF-bound Ctr-OCRs compared to all Ctr-OCRs. The circle size represents the percentage of either p63-bound Ctr-OCRs (left) or CTCF-bound Ctr-OCRs (right) in each CS. Relative score is calculated by comparing odds ratios between two overlap tests, in which odds ratio is estimated by BEDtools *fisher* option. C. Percentage of different loop types defined in the PCHiC data. D. Size distribution of promoter-enhancer loops in the PCHiC data. E. Estimated enrichment of CTCF and p63 at promoters, enhancers, or TAD Boundary compared to the reshuffled sequences (CTR). Odds ratio indicates the enrichment (> 1)/depletion (< 1) of two independent peak lists calculated by BEDtools *fisher* option.

Next, to analyze the localization of p63-bound and CTCF-bound Ctr-OCRs in regulatory chromatin interactions, we utilized the Promoter Capture Hi-C (PCHiC) data and TAD (topologically associating domain) regions from published HiC data in keratinocytes [18]. In the PCHiC, promoters were used as baits to capture the interacting genomic regions. In total, this PCHiC dataset captured 119,648 interactions, including 103,497 (86.5%) promoter-enhancer loops and 16,151 (13.5%) promoter-promoter loops (Figure 3C). As p63 plays a more prominent role at enhancers (Figure 3A) [4], we focused on promoter-enhancer loops. These loops had a median loop size of 250.9 kb (Figure 3D). Among the previously defined 1,223 p63-bound Ctr-OCRs (Figure 2C), 388 of them were located at the anchors of 1,674 loops, at either the promoter or the enhancer, and on average one p63-bound Ctr-OCR was connected by ∼4.3 loops. Similarly, among the 376 CTCF-bound Ctr-OCRs (Figure 2D), 191 of them were located at the anchors of 1,093 loops; on average, one CTCF-bound Ctr-OCR was connected by ∼5.7 loops. Therefore, CTCF-bound Ctr-OCRs at anchors were connected by more loops.

Furthermore, we found that CTCF-bound Ctr-OCRs located at loop anchors were more enriched for promoters (∼4 fold) (Figure 3E), consistent with all CTCF BSs and all CTCF Ctr-OCRs. In contrast, p63-bound Ctr-OCRs associated with loops were enriched in both promoters and enhancers to a similar extent (Figure 3E), which is different from all p63 BSs and all p63-bound Ctr-OCRs that were only enriched at enhancers (Figure 3B). We also examined whether p63-and CTCF-bound Ctr-OCRs were localized at the TAD boundaries. As expected, CTCF-bound Ctr-OCRs showed ∼2.5 fold enrichment for TAD boundaries, whereas p63-bound Ctr-OCRs were depleted of TAD boundaries (Figure 3E). Randomization by reshuffling genomic regions with same sizes did not show any enrichment in TAD boundaries, promoters or enhancers. As CTCF-bound Ctr-OCRs were enriched at the promoters and p63-bound Ctr-OCRs were enriched at both promoters and enhancers, it is plausible that p63-bound enhancers interact with CTCF-bound promoters to regulate genes through DNA looping and a subset of these loops may be affected by p63 mutations.

### p63 and CTCF mediate a subset of loops to regulate transcription

To assess how p63 and CTCF cooperates via DNA looping to regulate gene expression, we performed further analyses using previously published gene expression data [13] to examine the relation between gene expression and DNA looping. In control keratinocytes, we defined highly expressed genes by quartiles to four groups using their transcriptional levels (Q1-4, FPKM ≥ 1), and another group with lowly expressed genes (Q0, FPKM < 1). We detected an increasing number of associated loops with elevated gene expression levels (Figure 4A). This suggests that highly expressed genes may be more indispensable of regulation through loops, which is line with previous study showing multi-loop activation hubs at key regulatory genes [19]. Moreover, the differentially expressed genes between control and p63 mutant keratinocytes (Figure 1B) were associated with significantly more loops (*p* < 0.001) than all expressed genes (Figure 4B). We also performed the same analysis among loops with either p63-bound or CTCF-bound Ctr-OCRs. Similar results were found, although in these comparisons the difference was not significant, probably due to the low number of associated loops (Figures 4C and D).

**Figure 4.**
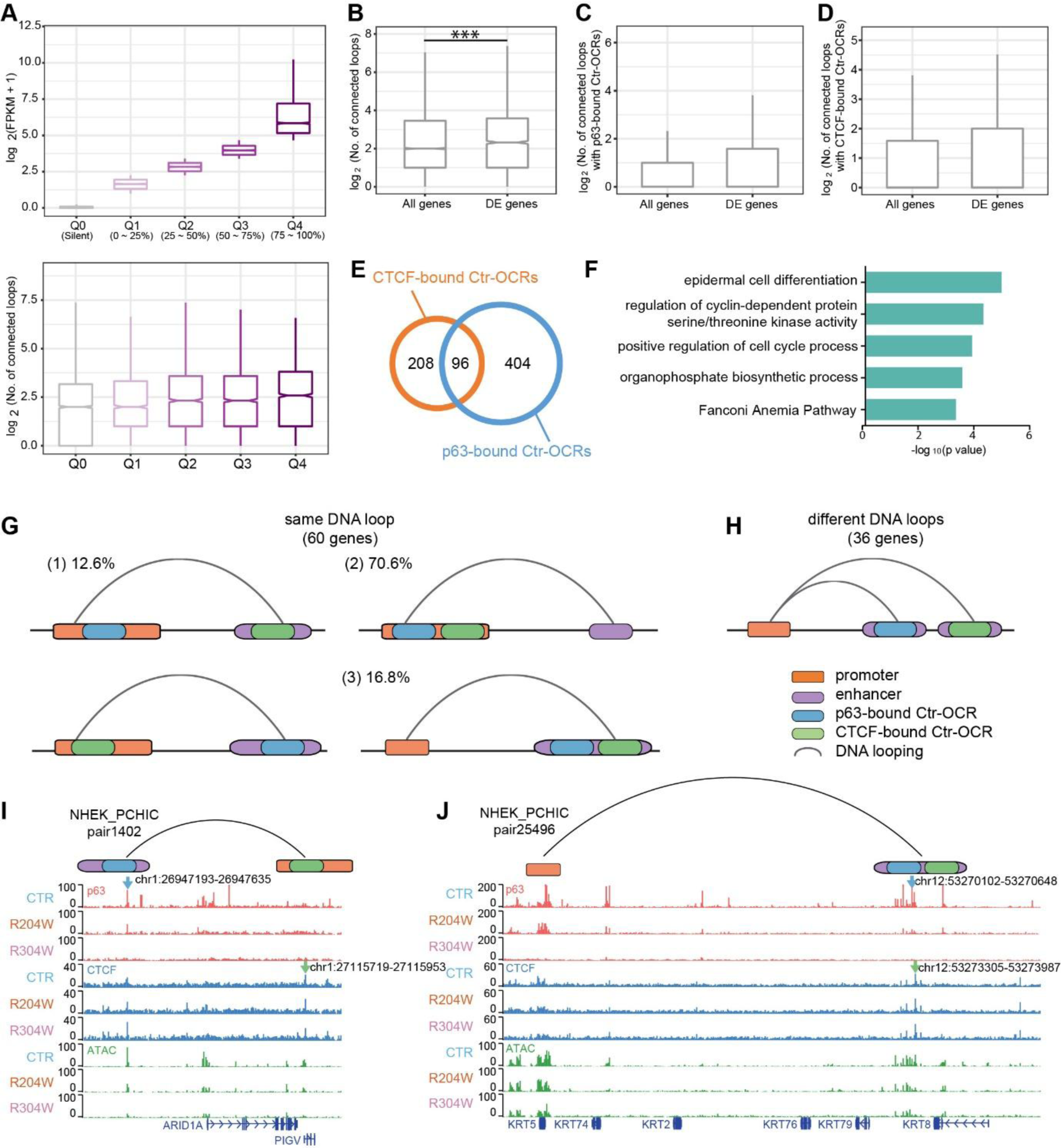
p63- and CTCF-bound Ctr-OCRs are involved in chromatin loops regulating epidermal genes. A. Highly expressed genes are associated with more DNA loops. Genes were divided in quarters according to their expression levels in keratinocytes. The number of connected DNA loops associated with each group of genes were plotted in the bottom bar chart. B. DE genes were associated with more connected DNA loops when compared to all expressed genes. ***, *p* < 0.001, Wilcoxon signed-rank test. C. DE genes were associated with more connected DNA loops with p63-bound Ctr-OCRs when compared to all expressed genes. D. DE genes were associated with more connected DNA loops with CTCF-bound Ctr-OCRs when compared to all expressed genes. E. Overlapping of genes mapped by CTCF-bound Ctr-OCRs (orange) and p63-bound Ctr-OCRs (blue). F. GO annotation of the 96 genes potentially co-regulated by p63 and CTCF. G. p63-bound and CTCF-bound Ctr-OCRs were involved in the same or different loops connecting the promoters of the 96 genes. 60 genes were regulated by the same loops with both p63-bound and CTCF-bound Ctr-OCRs: (1) in 12.6% cases, p63-bound Ctr-OCR located at one anchor while CTCF-bound Ctr-OCR located at the other anchor of the loop; in the majority cases, p63-bound and CTCF-bound Ctr-OCRs were both present at the (2) promoter (70.6%) or (3) enhancer (16.8%) of the DNA loops. H. The left 36 genes were co-regulated by connecting DNA loops with p63-bound and CTCF-bound Ctr-OCRs at enhancers of different loops. I. *PIGV* is shown as an example of G (1), which is connected by the DNA loop pair1402, bridging the PIGV promoter (with a CTCF-bound Ctr-OCR) and a distal enhancer (with a p63-bound Ctr-OCR). J. *KRT5* is shown as an example of G (3), which is connected by the DNA loop pair25496, bridging the KRT5 promoter and a distal enhancer (with both CTCF-bound and p63-bound Ctr-OCRs). The blue arrows pointed to the p63 BSs while the green arrows pointed to the CTCF BSs.

We then mapped 388 p63-bound and 191 CTCF-bound Ctr-OCRs which are located at loop anchors to their target genes by identifying their interacting gene promoters located at anchors of the other end of the loops. In some cases, Ctr-OCRs were located in promoters, and the associated genes were considered as target genes. In total, 388 p63-bound Ctr-OCRs interacted with 304 gene promoters while 191 CTCF-bound Ctr-OCRs interacted with 500 gene promoters through looping interactions, indicating that on average a CTCF-bound Ctr-OCR is involved in the regulation of more genes than a p63-bound Ctr-OCR. There was a significant (*p* < 0.001) overlap (N = 96) of genes that were potentially regulated by p63-and CTCF-bound Ctr-OCRs (Figure 4E). These 96 genes represent genes connected by the promoter-enhancer loops that were anchored by both p63-bound and CTCF-bound Ctr-OCRs and that may be affected in p63 mutant keratinocytes. As the three-dimensional chromatin landscape is relatively stable, we reasoned that the difference of the chromatin landscape in control and p63 mutant keratinocytes may influence gene expression during keratinocyte differentiation [18]. Therefore, we examined the expression of these 96 genes using previously reported differential gene expression in control and p63 mutant keratinocytes during differentiation [13]. Indeed we found that 39 of these 96 genes showed deregulation (*p* < 0.001). Notably many of the 96 genes were involved in ‘epidermal cell differentiation’, e.g. *KRT5, KRT75*, and *B9D1* (Figure 4F).

Next, we dissected how p63-bound and CTCF-bound Ctr-OCRs at loop anchors co-regulated these 96 genes: whether p63-bound and CTCF-bound Ctr-OCRs were involved in the same or different promoter-enhancer loops. We found that 60 genes were regulated by promoter-enhancer loops where p63-bound and CTCF-bound Ctr-OCRs were both located at the anchors in the same loops, although they may co-localized in different manner (Figure 4G). Among these 60 genes, 26 showed differential expression between control and p63 mutant keratinocytes during differentiation, with *PIGV* and *KRT5* shown as examples (Figures 4I and J). Among the loops associated with the 60 genes, the case where p63-bound Ctr-OCR was located at one anchor while CTCF-bound Ctr-OCR was located at the other anchor of the loop counted for 12.6% (Figure 4G, (1)). In the majority of the cases, both p63-bound and CTCF-bound Ctr-OCRs were present either at the promoter (70.6%) (Figure 4G, (2)) or at the enhancer (16.8%) (Figure 4G, (3)). Interestingly, for the rest 36 genes gene promoters were connected with both p63-bound and CTCF-bound Ctr-OCRs from different loops (Figure 4H), suggesting that enhancers occupied by p63-bound and CTCF-bound Ctr-OCRs can co-regulate the same gene through different loops. Among these 36 genes, 13 were deregulated in p63 mutant keratinocytes during differentiation.

## Discussion

Over the years, molecular understanding of p63 function in keratinocytes has led to the identification of numerous target genes including chromatin factors [20-22] as well as of a solid role of p63 in orchestrating the enhancer landscape [4, 5, 23]. However, whether p63 is involved in the direct regulation of chromatin architecture is not known. In this study, we characterized the chromatin accessibility with ATAC-seq in both control and EEC syndrome patient keratinocytes carrying p63 mutations (R204W and R304W). Interestingly, we found decreased chromatin accessibility in p63-and CTCF-bound open chromatin regions in mutant p63 keratinocytes. These less accessible chromatin regions interact with epidermal gene promoters and potentially regulate gene expression. Collectively, we proposed an unreported role of p63 directly cooperating with CTCF in chromatin interactions.

It is envisaged that p63 is a key factor in skin keratinocytes [4, 5, 24]. Ample studies have shown that the proper regulation of gene expression by p63 through the precise control of enhancers is essential for maintaining the epidermal cell identity [4, 13, 25, 26]. It is equally important that p63 requires additional co-regulating TFs to regulate transcription. Several studies reported p63 co-regulating TFs, e.g. AP1, AP2, STAT5, and RFX5 [20, 22, 23, 27], which are essential for the epidermal gene expression in a temporal and spatial manner [4, 28]. Besides, p63 interacts with the chromatin remodelling factor BAF1 to maintain the open chromatin regions [5] and with Dnmt3a to locate enhancers [12]. In this study, we found that CTCF, a well-known TF maintaining the chromatin architecture, is also a p63 co-regulating TF modulating a subset of chromatin loops. However, this cooperation is probably not through direction protein-protein interactions but mainly bridging enhancers and promoters, given the few overlap observed between p63 and CTCF BSs. It is known that the majority of p63-bound regions are located in intergenic or intragenic regions (enhancers) rather than promoters [22], and therefore how p63-bound enhancers interact with gene promoters is an interesting question. Our current study provides the first line of evidence that the cooperation with CTCF assists the looping of p63-bound enhancers to gene promoters.

The observation of the enriched CTCF motif in Ctr-OCRs detected in our ATAC-seq analyses was somewhat unexpected. Our previous enhancer-centred analysis that focused on H3K27ac enriched genomic regions did not capture CTCF motifs [4, 13], probably due to the fact that CTCF is more enriched in open chromatin regions depleted of H3K27ac but enriched for H3K4me3 signal (Figure 3A and Supplementary Figure 3). Interestingly, detecting the CTCF motif in Ctr-OCRs in keratinocytes is in line with the previous finding that the motif of YY1 was enriched in p63-bound sequences [27]. YY1 has been shown to co-localize with CTCF and stabilize chromatin looping [29, 30].

CTCF maintains the genome organization by occupying the boundaries of megabase-scale TADs, and functions as an insulator to block enhancer-promoter interactions between different TADs [16]. Our date showed that, in addition to demarcating TADs, CTCF mediates promoter-enhancer loops, often located in promoter-proximal regions (Figure 3E), to facilitate the promoter-enhancer interactions within one TAD. This is in line with the concept that a subpopulation of CTCF associates with the RNA polymerase II (Pol II) protein complex to activate transcription [31]. It is likely that CTCF helps to bridge the p63-bound enhancers to transcription start site-proximal regulatory elements and to initiate transcription by interacting with Pol II, thus supporting a role of CTCF in facilitating contacts between transcription regulatory sequences. This model has been demonstrated by the previous work on the beta-globin locus [32]. To get a clearer picture, technologies that are similar to ChIA-PET [33], such as HiChIP [34], would be useful to understand long-range contacts associated with CTCF or p63 in skin keratinocytes.

The three-dimensional chromatin landscape is relatively stable once established in a specific cell type, probably during cell commitment, and cell-type-specific looping structures mainly control the accessibility of enhancers to their specific targets [35]. Recent studies showed that cell type-specific TFs are involved in regulation of DNA looping in macrophage development, which shed light on the role of cell type-specific TFs in regulating chromatin interactions [19]. Here we proposed that a number of loci nearby epidermal genes were organized into a ‘regulatory chromatin hub’ within the chromatin interactions mediated by CTCF in epidermal keratinocytes (Figure 5A). Such hubs contain multiple connecting DNA loops that require not only CTCF binding that is rather static but also binding of cell type-specific TFs (e.g. p63) for the transcriptional activity. In this model, cell type-specific TFs may be essential to make the DNA loops active in transcription. This hypothesis is consistent with our observation that unchanged CTCF binding signal but decreased accessibility in the CTCF-bound Ctr-OCRs in p63 mutant keratinocytes (Figure 2D).

**Figure 5.**
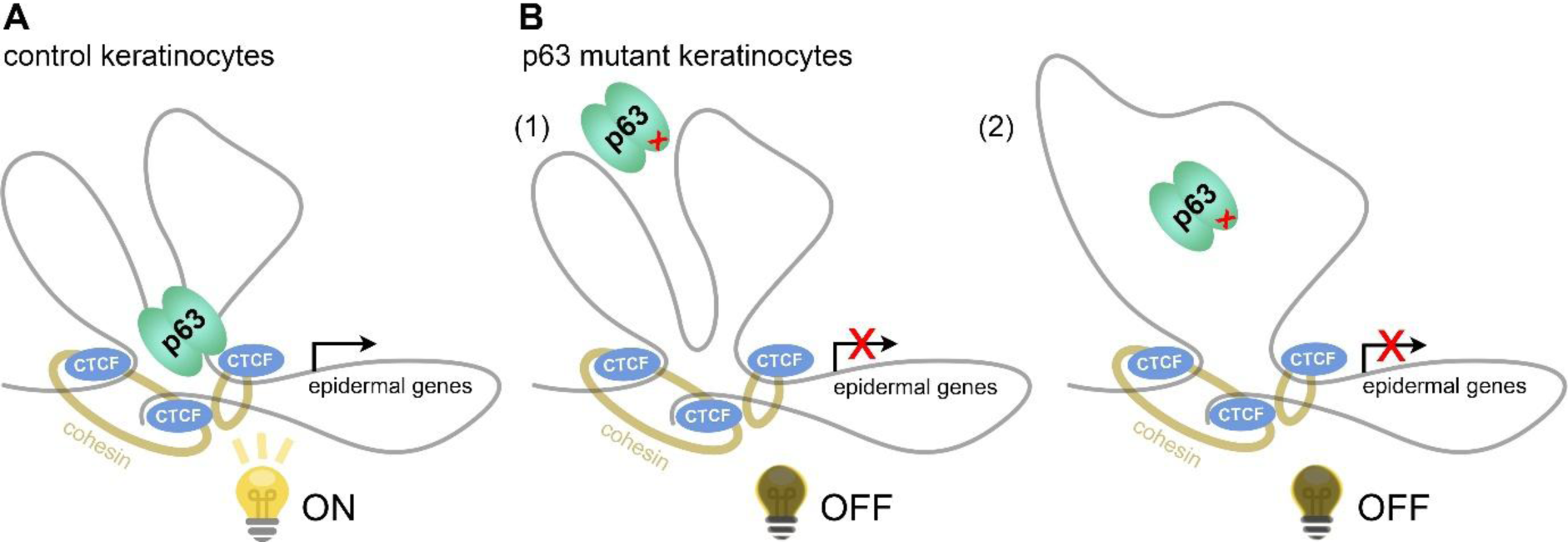
Models of how p63 and CTCF co-mediate chromatin loops regulating epidermal genes. A. In control keratinocytes, p63 and CTCF co-mediate a subset of DNA loops nearby epidermal genes and regulate their gene expression. B. In p63 mutant keratinocytes, due to the loss of p63 binding, p63 and CTCF co-mediated DNA loops are not functional anymore, and therefore epidermal genes are deregulated. There are two possibilities in deregulated function of looping: (1) the looping structure is stable as CTCF binding is unchanged; (2) the looping structure is partially changed because of loss of p63 binding and loss of the p63-dependent loop.

Moreover, our observations shed light on the chromatin-based mechanisms underlying EEC syndrome caused p63 mutations. These EEC mutations, such as R204W and R304W, were shown to disrupt p63 DNA binding, resulting in impaired transactivation activity and loss of epidermal cell identity [13, 36, 37]. Our data suggest that deregulated function of DNA loops mediated by p63 and CTCF represents an additional layer to the disease mechanism (Figure 5B). Given that CTCF binding did not change, it is likely that the looping interactions are static, but the looping is no longer active due to the absence of p63 binding (Figure 5B (1)). Therefore, the epidermal genes were down-regulated and p63 mutant keratinocytes showed loss of epidermal cell identity [13]. On the other hand, it is also possible that the loop ablation as a result of lost p63 binding leads to the decreased accessibility at CTCF-bound promoters and deregulated gene expression in p63 mutant keratinocytes (Figure 5B (2)). The latter scenario suggests p63 is the key factor determining the regulatory activity of such promoter-enhancer interactions. Nevertheless, how p63 mutations affect the cooperation between p63 and CTCF and gene expression is still unclear. Among the 96 genes with loops connected by p63-and CTCF-bound Ctr-OCRs, only 39 of them were deregulated in p63 mutant keratinocytes. It is possible that ‘shadow enhancers’ that are composed of clusters of enhancer and contribute to robust gene transcription [38, 39], are involved in regulation of the 57 genes that are not deregulated in mutant keratinocytes. Further experiments are required to test these hypotheses.

Taken together, our results highlight the cooperation of p63 and CTCF in mediating DNA loops as well as gene regulation. These observations provide new insights into the general regulatory role of chromatin looping which depends on the coordination of CTCF and cell type-specific TFs such as p63 in skin keratinocytes.

## Material and methods

### Ethics statement

All procedures for establishing and maintaining human primary keratinocytes were approved by the ethical committee of the Radboud university medical center (“Commissie Mensgebonden Onderzoek Arnhem-Nijmegen”). Informed consent was obtained from all donors of a skin biopsy.

### Human primary keratinocyte culture

Primary keratinocytes were established previously from skin biopsies of three EEC syndrome patients carrying heterozygous mutations in the p63 DNA-binding domain, R204W[40] and R304W [37], as well as of non-EEC volunteers (Dombi23, referred to as control) [41]. R204W and R304W are treated as a p63 mutant group for further analyses to minimize the effect from individual difference. As previously described [42], primary keratinocytes were cultured in Keratinocyte Basal Medium (KBM, Lonza #CC-4131) supplemented with 100 U/mL Penicillin/Streptomycin (Gibco Life Technology #15140122), 0.1 mM ethanolamine (Sigma Aldrich #141-43-5), 0.1 mM O-phosphoethanolamine (Sigma Aldrich #1071-23-4), 0.4% (vol/vol) bovine pituitary extract, 0.5 μg/mL hydrocortisone, 5 μg/mL insulin and 10 ng/mL epidermal growth factor (Lonza #CC-4131). Medium was refreshed every other day. When cells were more than 90% confluent, cells were collected at the proliferation stage. No mycoplasma contamination is found during cell culture.

### ATAC-seq

ATAC libraries of the control and p63 mutant keratinocytes were prepared by a documented protocol [43]. In brief, keratinocytes were treated with Accutase® solution on plates and well re-suspended into single cells. Pellet cells for 5 min at 500 g. After twice wash with ice-old PBS, pellet the cells again. Re-suspend cells and take out 100,000 cells for lysis with ice-old freshly made lysis buffer (containing 10 mM Tris-HCl pH 7.5, 10 mM NaCl, 3 mM MgCl_2_ and 0.1% IGEPAL CA-630 detergent). Then perform tagmentation using 2 μl of Tn5 transposase and 12.5 ul 2 × TD buffer (Illumina #FC-121-1031) at 37°C for 1h with 650 rpm shaking. The resulted DNA fragments underwent two sequential seven-cycle PCR amplification, and in between the libraries were selected for <500 bp fragments using SPRI beads. The final PCR products were purified with QIAquick PCR Purification Kit (QIAGEN #28106) and quantified with the KAPA Library Quantification Kit (Kapa Biosystems #KK4844), and then sequenced in a paired-ended manner using the NextSeq 500 (Illumina) according to standard Illumina protocols.

### CTCF ChIP-seq

Chromatin for ChIP was prepared as previously described [20]. ChIP assays were performed following a standard protocol [44] with minor modifications. On average, 0.5M keratinocytes were used in each ChIP. 4x ChIP reactions are pooled to prepare one ChIP-seq sample. Antibodies against CTCF (Millipore #07-729, 5ul) was used in each ChIP assay. Resulted DNA fragments from four independent ChIP assays were purified with QIAquick PCR Purification Kit (QIAGEN #28106). Afterwards, 5 ng DNA fragments were pooled and proceeded on with library construction using KAPA Hyper Prep Kit (Kapa Biosystems #KK8504) according to the standard protocol. The prepared libraries were then sequenced using the NextSeq 500 (Illumina) according to standard Illumina protocols.

### RNA-seq data analysis

Paired-end RNA-seq reads were mapped to the human genome hg19 using STAR aligner[45] in two-pass mode with default parameters, and enumerate stranded gene-level read counts at the same time. The generated count matrix was used as input for DESeq2 package [46] to distinguish differential expressed genes between control and mutant keratinocytes. These genes greater than 1.5-fold changed at adjusted *p* value < 0.05 were considered significantly deregulated.

### ChIP-seq and ATAC-seq data analysis

Sequenced reads were aligned against the UCSC hg19 human reference genome with Burrows-Wheeler Aligner (BWA) program with default parameters [47]. The potential PCR and optical duplicates were removed using Picard *MarkDuplicates* option. The filtered BAM files were inputted to MACS2 [48] for peak calling. The p63 ChIP-seq data was published and re-analyzed in this study [13]. The p63, CTCF and H3K27ac were called using the narrow setting (default) with a q-value of 0.01. For ATAC-seq, the open chromatin regions were predicted using default parameters except for using -f BAMPE option. Peaks overlapping with the consensus excludable ENCODE blacklist were dropped to avoid confounding by repetitive regions. All alignment files were extended to 200-bp and scaled to RPKM-normalized read coverage files using deepTools [49] for visualization. To compare binding profiles between different samples unbiasedly, we applied library size factors estimated from DESeq2 [46] on RPKM values.

Differentially accessible regions were detected using DESeq2 package [46] with fold change less than 2.0 and p-value below 0.05. Differential motif analysis in the differential DHSs was employed by the findMotifs function in HOMER tool (http://homer.salk.edu/homer/motif/) with other default parameters, which can normalize the background sequences to remove GC-bias. The BEDtools suite (https://bedtools.readthedocs.io/en/latest/content/bedtools-suite.html) was used to test overlap and enrichment between different intervals.

### Dimensionality reduction and functional annotation

For visualization, top 1,000 variable genes were first selected based on interquartile range (IQR) of normalized expression values, and further used to reduce dimensionality (principal component analysis) of the dataset by pca function in R. Functional enrichment was evaluated by Metascape online tool [50], to gain insight into the biological functions for deregulated genes. Only these functional terms with Benjamini-adjusted p-value < 0.05 were considered significantly overrepresented.

### Capture Hi-C data processing

Raw Promoter Capture Hi-C sequencing reads from GSE84662 were processed using the HiCUP pipeline [51], like quality control, alignment to hg19 and reads filtering. Technical replicates were merged and de-duplicated, and then pooled biological replicates was then jointly used for interaction identification with CHiCAGO [52] and the associated chicagoTools suite. All significant interactions were defined based on a strict interaction threshold (CHiCAGO score ≥ 5).

Briefly, CHiCAGO predicts interactions based on a convolution background model reflecting both ‘Brownian’ (real, but expected interactions) and ‘technical’ (assay and sequencing artifacts) components. The putative p values are corrected using a weighted false discovery control procedure that specifically accommodates the fact that increasingly larger numbers of tests are performed at regions where progressively smaller numbers of interactions are expected. The weights were learned based on the decrease of the reproducibility of interaction calls between the individual replicates of macrophage samples with distance. Interaction scores were then computed for each fragment pair as –log-transformed, soft-thresholded, weighted p values.

## Author Contributions

JQ, GY and HZ conceived and designed the experiments. JQ, GY, HZ wrote and revised the manuscript. JQ performed the experiments. JQ, GY and HZ analysed the data.

## Acknowledgements

We thank Eva Janssen-Megens, Siebe van Genesen and Rita Bylsma for operating the Illumina analyzer. We thank the ENCODE Consortium for sharing their data.

## Competing interests

None of the authors have any competing interests in the manuscript.

## Data availability

To review our complete dataset GEO accession GSE123711 (https://www.ncbi.nlm.nih.gov/geo/query/acc.cgi?acc=GSE123711). All data supporting the findings of the study and in-house codes are available on request.

## Ethics approval and consent to participate

Not applicable.

## Declarations

### Consent for publication

Not applicable.

### Funding

This research was supported by Netherlands Organisation for Scientific Research (NWO/ALW/MEERVOUD/836.12.010, HZ), Radboud University fellowship (HZ) and Chinese Scholarship Council grant 201406330059 (JQ).

## Supplementary Figures

**Supplementary Figure 1.**
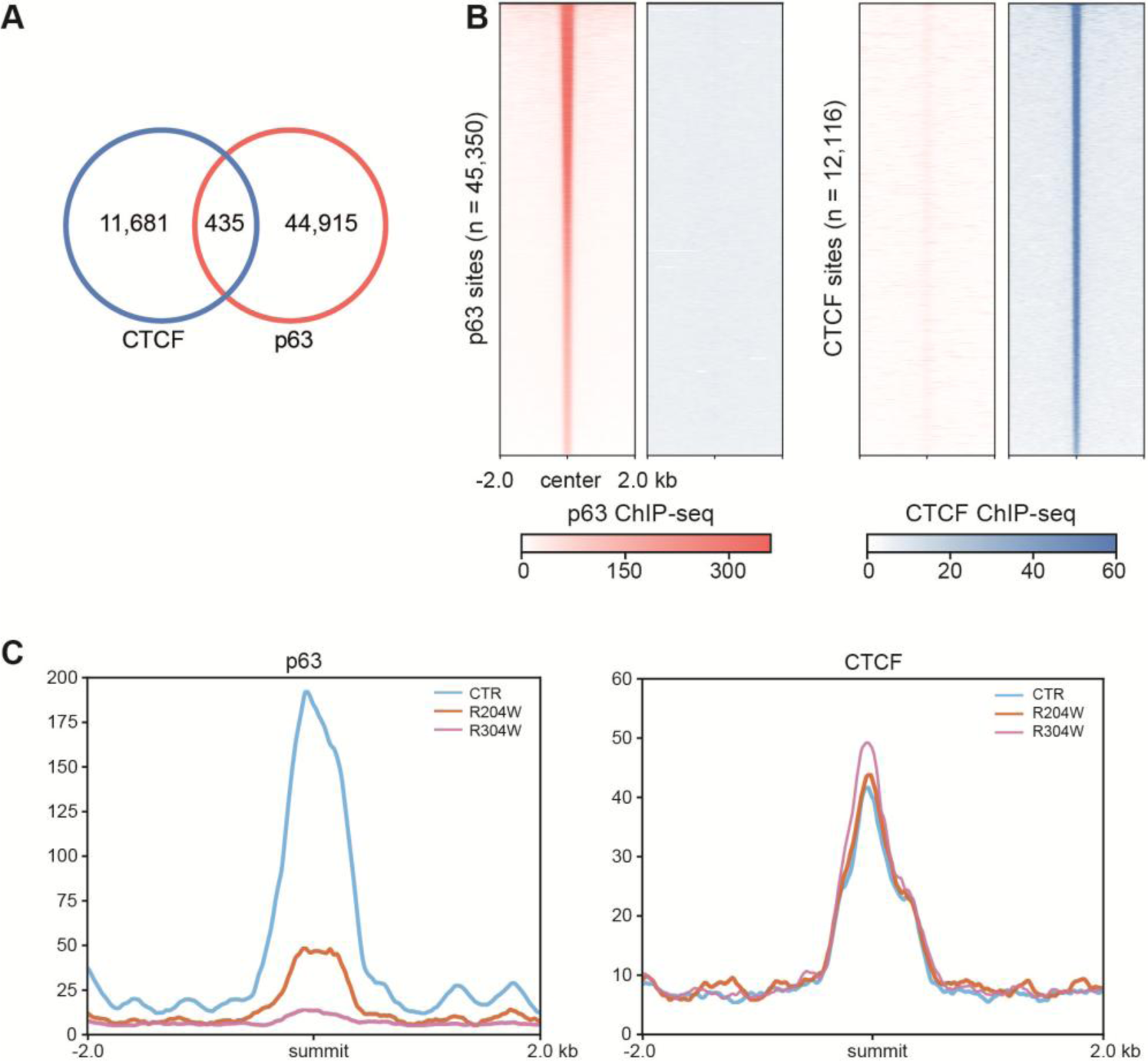
Few overlap between p63 and CTCF binding sites. A. Overlapping of p63 and CTCF binding sites from ChIP-seq data B. Heatmaps showing the CTCF binding signal at all 45,350 p63 binding sites (left) and the p63 binding signal at all 12,116 CTCF binding sites (right). C. Bandplots showing the p63 binding signal (left) and CTCF binding signal (right) at the 435 co-binding sites in both control and p63 mutant keratinocytes.

**Supplementary Figure 2.**
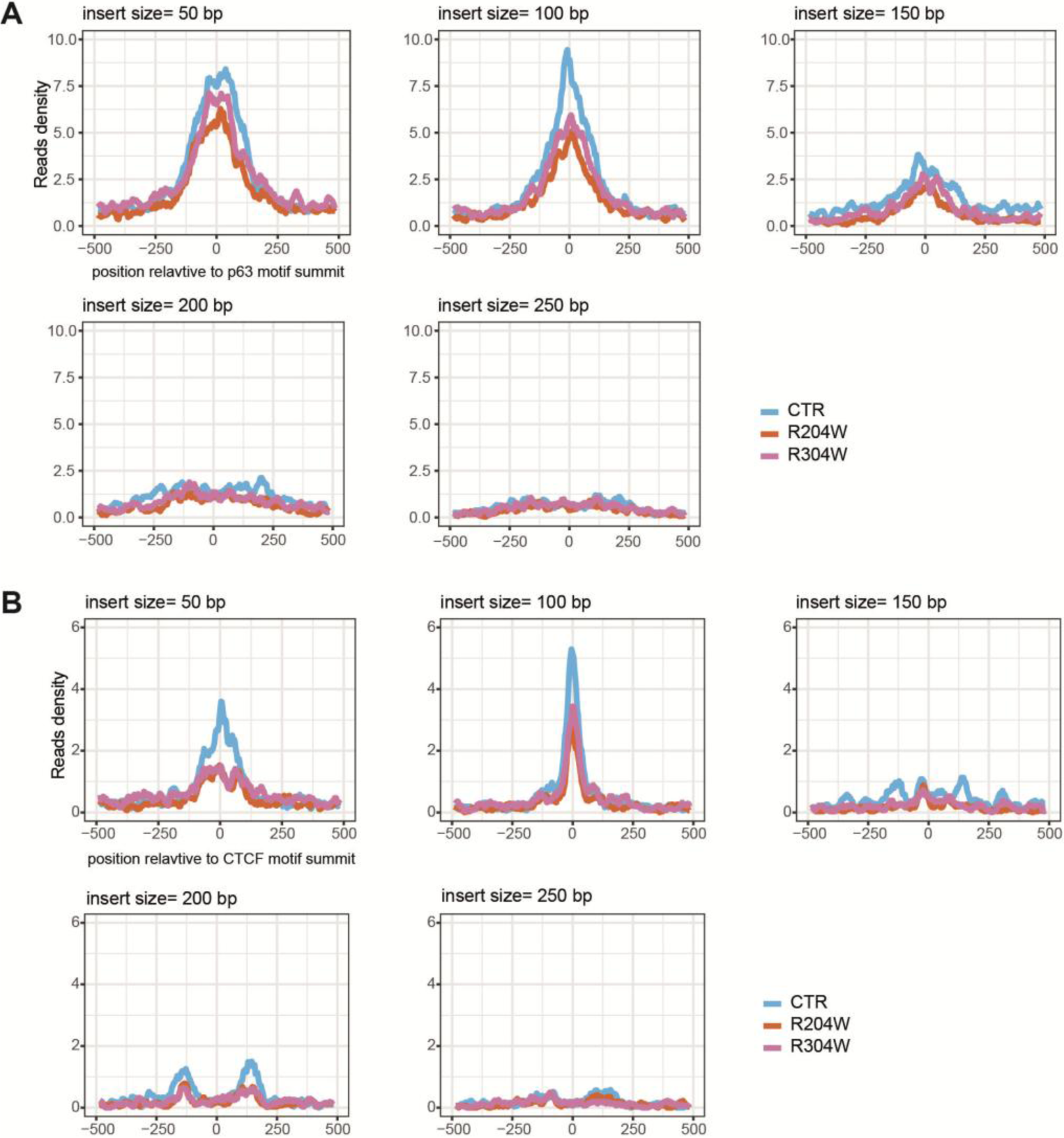
Deregulated nucleosome organization at p63-bound and CTCF-bound Ctr-OCRs in p63 mutant keratinocytes. D. Reads density of ATAC-seq at p63-bound Ctr-OCRs with different insert sizes in both control and p63 mutant keratinocytes. E. Reads density of ATAC-seq at CTCF-bound Ctr-OCRs with different insert sizes in both control and p63 mutant keratinocytes.

**Supplementary Figure 3.**
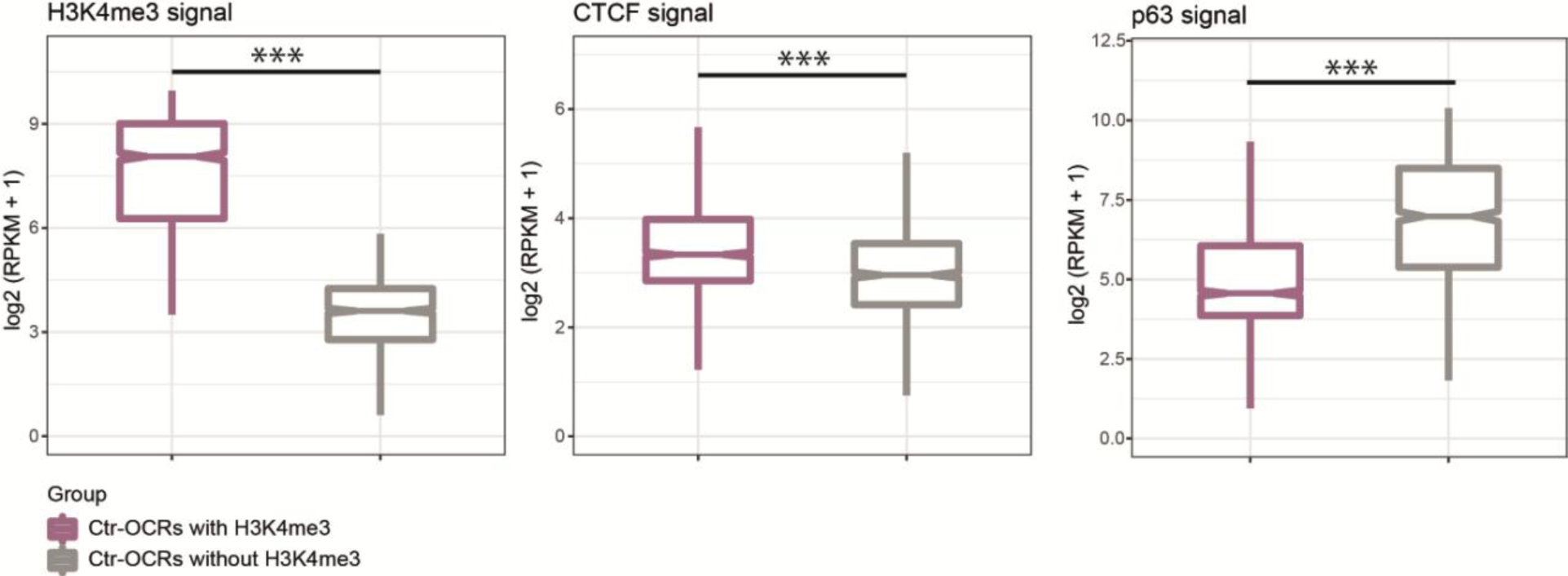
CTCF bound preferentially to Ctr-OCRs marked by H3K4me3. Ctr-OCRs are grouped into two categories according to H3K4me3 signal, Ctr-OCRs with H3K4me3 (purple) and Ctr-OCRs without H3K4me3 (gray). CTCF ChIP-seq showed higher signal in Ctr-OCRs with H3K4me3 while p63 ChIP-seq showed higher signal in Ctr-OCRs without H3K4me3 in control keratinocytes. ***, *p* < 0.001, Wilcoxon signed-rank test.

